# Construction of Nanobody Library in Mammalian Cells by Linear-double-stranded DNA Based AND Gate Genetic Circuit

**DOI:** 10.1101/2020.01.21.912907

**Authors:** Yanjie Zhao, Xin Tong, Chunze Zhang, Shuai Li

## Abstract

Nanobody is one special type of single-domain antibody fragment with multiple advantages over traditional antibody. Our previous work established linear-double-stranded DNA (ldsDNA, or PCR amplicon) as novel biological parts for building AND gate genetic circuits in mammalian cells. During this AND-gate circuit formation process, the co-transfected up- and down-stream ldsDNAs could be linked together to form intact gene expression cassette. Here, we employed this **<u>l</u>**dsDNA-**<u>b</u>**ased **<u>A</u>**ND-**<u>g</u>**ate (LBAG) strategy to construct nanobody library in mammalian cells. The sequence complexity of complementary determining regions (CDRs) was introduced into ldsDNA by PCR amplification. After being co-transfected into mammalian cells, the up- and down-stream ldsDNAs undergo AND gate linkage and form full nanobody coding regions, containing CDR1-3. High throughput sequencing identified 22,173 unique oligonucleotide sequences in total generated by this strategy. Thus, we developed a novel method to construct nanobody library, which is a start point for building high content nanobody library in mammalian cells.

## Introduction

Nanobody, also known as single-domain antibody (sdAb) or domain antibody, is one special type of antibody consisting of a single monomeric variable antibody domain (1,2). Since being first discovered in camelids in 1993, nanobody has been serving as a good platform for the development of therapeutic antibody due to its advantages such as small size (∼15 kDa, 4 nm long and 2.5 nm wide), high stability, high solubility and excellent tissue penetration ability (2-6). In 2019, the first nanobody-based medicine, Cablivi® (caplacizumab), was approved by FDA for the treatment of acquired thrombotic thrombocytopenic purpura (aTTP) (7). Meanwhile, multiple other nanobodies are currently under clinical development (7,8).

Different state-of-the-art display technologies, e.g., phage, bacteria, yeast, ribosome, and mRNA-display, are used to identify candidate nanobodies from large-size nanobody libraries (9-15). These technologies associate the phenotype (affinity between nanobody and antigen) to the genotype (DNA sequence of nanobody). After rounds of selection, by Sanger or high throughput sequencing, the sequences encoding high affinity nanobodies can be obtained (5). However, in current display technologies, nanobodies are generated and displayed in non-mammalian systems, which limits their applications. For example, it’s hard to express and screen transmembrane proteins, such as G protein-coupled receptors (GPCRs), ion-channels, which are the most attractive therapeutic target classes (16,17). Although several different approaches, e.g. nanodisc (18,19), next-generation detergent formulation (20,21), have been developed to facilitate the screen of transmembrane proteins, the ideal platform would allow direct panning of display libraries against intact mammalian cells with functional signal transduction pathways (22-24).

Mammalian cells are good platforms for antibody development, as the appropriate protein chemical modification, and natural folding structure of the transmembrane proteins. Ingenious design methods, such as mammalian cell surface display, and autocrine-based signaling system, have been developed to facilitate the panning of antibody in human cell lines (22,23,25-27). Currently, immune-derived library or focused-phage library were applied to cultured cells to select candidate hits (22,28,29). However, for synthetic plasmid-based library, the main bottleneck is the achievable library size (10^9^), which is limited by transformation efficiency. Moreover, when introduce into mammalian cells, the complexity of synthetic library keeps on losing during plasmid preparation and cell-transfection process. So, it’s still hard to introduce high content synthetic nanobody library into cultured mammalian cells.

linear-double-stranded DNA (ldsDNA, or PCR amplicon) is a good medium to incorporate and store highly complex genetic coding information. Our previous research demonstrated that ldsDNA could relink together to form AND gate genetic circuits in mammalian cells (30). We started to construct DNA-encode library by this ldsDNA-based AND-gate strategy. Indeed, we employed terminal-NNKs-ldsDNA-based AND-gate genetic circuit to generate high content peptide library (31). In this study, we developed a proof-of-concept **<u>l</u>**dsDNA-**<u>b</u>**ased **<u>A</u>**ND-**<u>g</u>**ate (LBAG) strategy to construct nanobody library in mammalian cells.

## Materials and Methods

The following protocols are basically the same as our previous methods with some modifications (30,31).

### Cell culture

HEK293T cell line was maintained in Dulbecco’s modified Eagle medium (DMEM) (Thermo Fisher) containing 10% fetal bovine serum (FBS) and 1% penicillin-streptomycin solution at 37°C with 5% CO_2_.

### ldsDNA synthesis

Up- and down-stream template ldsDNAs were synthesized by GENEWIZ (Suzhou, China) and then subcloned into pUC57 vector, respectively (for template ldsDNA sequences see Supplementary Materials). KOD DNA polymerase (Toyobo) was used to generate up- and down-stream ldsDNAs (PCR amplicons) taking up- or down-stream template ldsDNA containing pUC57 plasmids as templates, respectively. To remove plasmid template and free dNTPs, PCR products underwent agarose electrophoresis and gel-purification (Universal DNA purification kit, Tiangen). The amount of PCR products was determined by OD 260 absorption value. PCR primer sequences were present in Supplementary Table 1.

### ldsDNA transfection

HEK293T cells were seeded into 6-well plates the day before transfection (60% – 70% confluency). 500 ng/well of each input amplicons (1000ng/well in total) were co-transfected into HEK293T cells with Lipofectamine 2000 (Invitrogen) reagent following standard protocol.

### RNA extraction, cDNA synthesis and PCR amplification

48 h after transfection, total RNAs were extracted with RNAiso Plus (TaKaRa) reagent. cDNA was synthesized by PrimeScript ™ RT reagent Kit with gDNA Eraser (TaKaRa) following standard protocol. KOD DNA polymerase was used to amplify the sequences flanking the ldsDNA junctions. PCR primers were present in Supplementary Table 1.

### Library construction and high-throughput sequencing

Library construction and high-throughput sequencing were conducted by Novogene (Tianjin, China). Sequencing libraries were generated using TruSeq® DNA PCR-Free Sample Preparation Kit (Illumina) following the manufacturer’s recommendations, and the index codes were added. The library quality was assessed on the Qubit Fluorometer and Agilent Bioanalyzer 2100 system. At last, the library was sequenced on an Illumina NovaSeq 6000 system and 250-bp paired-end (PE) reads were generated.

### Data analysis

Paired-end reads were assigned to each sample according to the unique barcodes. Reads were filtered by QIIME quality filters. Paired-end reads were merged using FLASH (v1.2.11, default parameter). Nucleotide sequences generated by ldsDNA-based AND-gate were extracted by in-house Perl scripts.

## Results

### Rationale of nanobody library construction by <u>l</u>dsDNA-<u>b</u>ased <u>A</u>ND-<u>g</u>ate (LBAG) genetic circuit

In this study, we take a GFP-targeted nanobody as the framework of nanobody library (32-34). This GFP-nanobody (cAbGFP4, for amino acid sequence see Supplementary Materials) has a very low Kd value (0.32 nM) and has already been unitized to isolate or track GFP-tagged proteins (34-36). Here, the CDR1 (containing 8 amino acids, 8AA), CDR2 (9AA) and CDR3 (7AA) of GFP-nanobody are replaced by the corresponding number of NNK degenerate codons (N = T/C/A/G, K = T/G) (33).

For the synthesis of upstream ldsDNAs, we designed a 132-bp-long 3’ primer, which contains 8 and 9 consecutive MNN codons (reverse complementary form of the NNK codons), corresponding to CDR1 and CDR2 respectively (Supplementary Table 1). There are a total of 34 Ns and 17 Ms in this 3’ primer. And the diversity of oligonucleotide sequence increases by three times with every N (T/C/A/G) nucleotide and increases by one time with every M (C/A) nucleotide. Thus, we get an extremely highly diverse 3’ primer pool in this way. Then we performed PCR amplification to introduce random CDR1 and CDR2 into the upstream ldsDNAs. In the primer annealing step of PCR reaction, the 3’ primers anneal to the template via its 3’ end 21-bp fragment (Figure 1A). And in extending step, DNA polymerase synthesizes the complementary strand using primer’s 5’ 111-bp fragment as template (Figure 1A). Therefore, the CDR1 and CDR2 sequence of every newly synthesized PCR product comes from the 3’ primer, thus it would be different from that of its template ldsDNA. After rounds of PCR cycles, we get an extremely highly diverse upstream ldsDNA pool, which contains nearly random CDR1 and CDR2.

**Figure 1.**
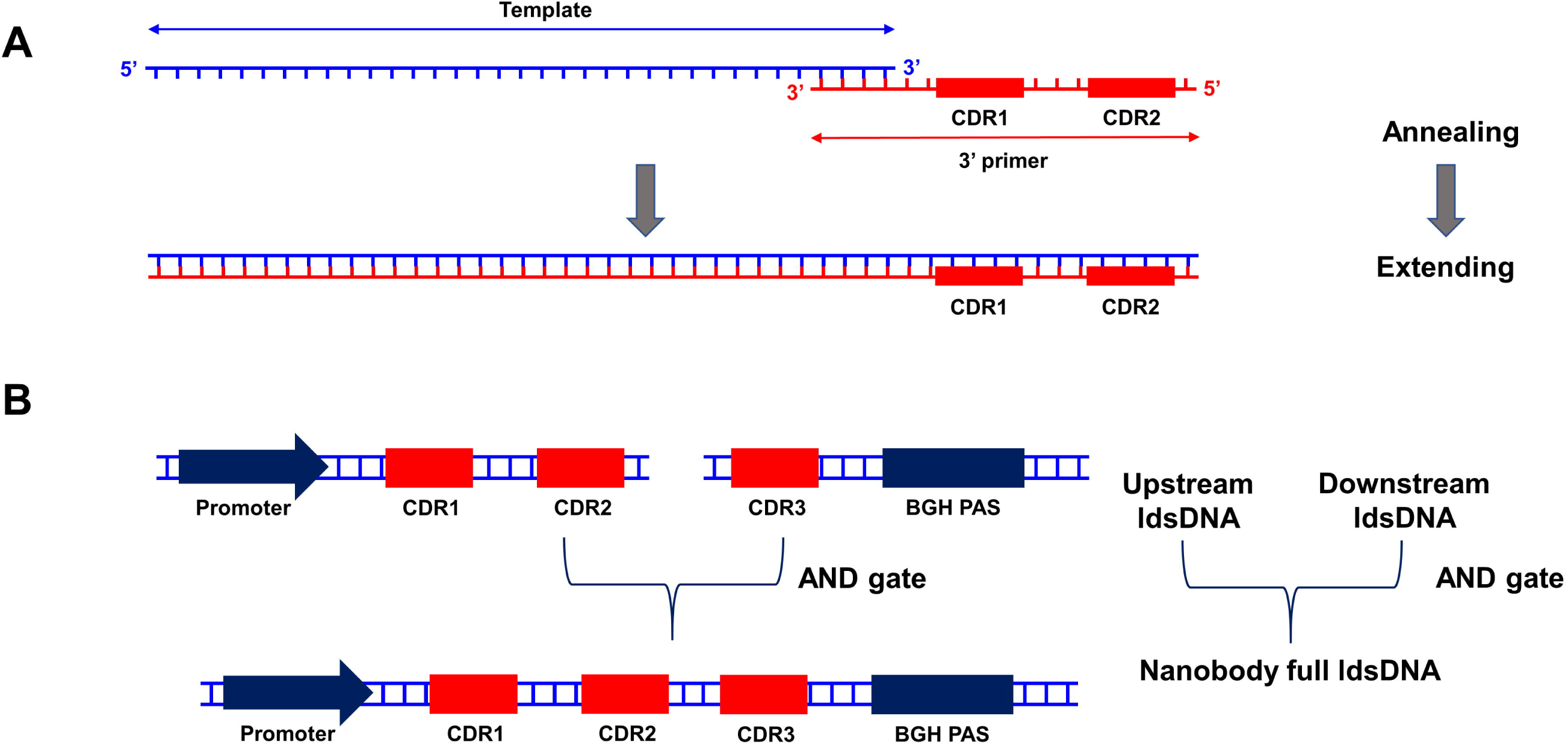
Schematic diagram of the generation of ldsDNA and the formation of LBAG genetic circuit. (A) Schematic showing the rationale of introducing CDR1 and CDR2 random sequence into up-stream ldsDNA by PCR amplification. In the primer annealing step, the 3’ primer binds to single-stranded template DNA through its 3’ fragment (21-bp). In the extending step, DNA polymerase would synthesize a complementary strand by taking the 5’ fragment of 3’ primer as template. Thereby, CDR1 and CDR2 sequences are incorporated into up-stream ldsDNAs. (B) In HEK293T cells, the up- and down-stream ldsDNAs link together through AND-gate calculation to form nanobody library generating full-length ldsDNAs.

For downstream ldsDNA, we designed a 143-bp-long 5’ primer, which contains 7 consecutive NNK codons corresponding to GFP-nanobody CDR3 (Supplementary Table 1). Similar to the synthesis process of upstream ldsDNAs, by PCR reaction, we get a highly diverse downstream ldsDNA pool, which contains random CDR3.

After being co-transfected into HEK293T cells, the upstream- and downstream-ldsDNAs link together through non-homologous end joining (NHEJ) mechanism and form full-length nanobody expression cassette following the AND gate principle (Figure 1B). The full-length cassette contains CMV promoter, CDR1, CDR2 sequences from upstream ldsDNA and CDR3, BGH poly A signal sequences from downstream ldsDNA. The CMV promoter would start nanobody mRNA transcription. The nanobody mRNAs would be translated into nanobody proteins in HEK293T cells. In this way, we construct nanobody library in mammalian cells.

### High throughput sequencing data statistics of AND gate nanobody library

Forty-eight hours after the transfection of ldsDNAs, total RNAs were extracted from HEK293T cells, cDNAs were synthesized, and PCR was performed to amplify CDR1-3 sequence formed by AND gate genetic circuit. CDR1-3 PCR amplicons were subjected to high throughput sequencing (HTS, NovaSeq 6000 PE250). Three biological repeats were performed. Raw PE250 HTS reads were quality filtered and merged. We combined clean merged paired-end reads from three biological repeats and got 4,173,356 reads. About 88.18% of merged reads (3,679,989) contain both upstream- and downstream-ldsDNA sequence and were subjected to further investigation.

### Length distribution of nanobody ldsDNA sequences formed by LBAG genetic circuit

In the NHEJ pathway, genomic dsDNA ends are subject to terminal processing, leading to either nucleotide(s) deletion or insertion (37). Our previous research also revealed that ldsDNAs transfected into HEK293T cells undergo end nucleotide(s) deletion (30). To evaluate ldsDNA ends processing during AND gate formation, we analyzed the length distribution of nanobody ldsDNAs. If terminal deletion or insertion happens to input ldsDNA, the length of full nanobody ldsDNA would change. As shown in Figure 2, the most abundant read length is 264-bp (154,477 reads), which is corresponding to the intact sequence length (implying that neither deletion nor insertion happens to ldsDNA ends). That is these most abundant 264-bp reads followed the same framework with GFP-nanobody (for sequence see Supplementary Materials). Other top read length is shorter than 264-bp (Figure 2). Thus, we can conclude that ldsDNA terminals usually undergo end nucleotide(s) deletion, not insertion.

**Figure 2.**
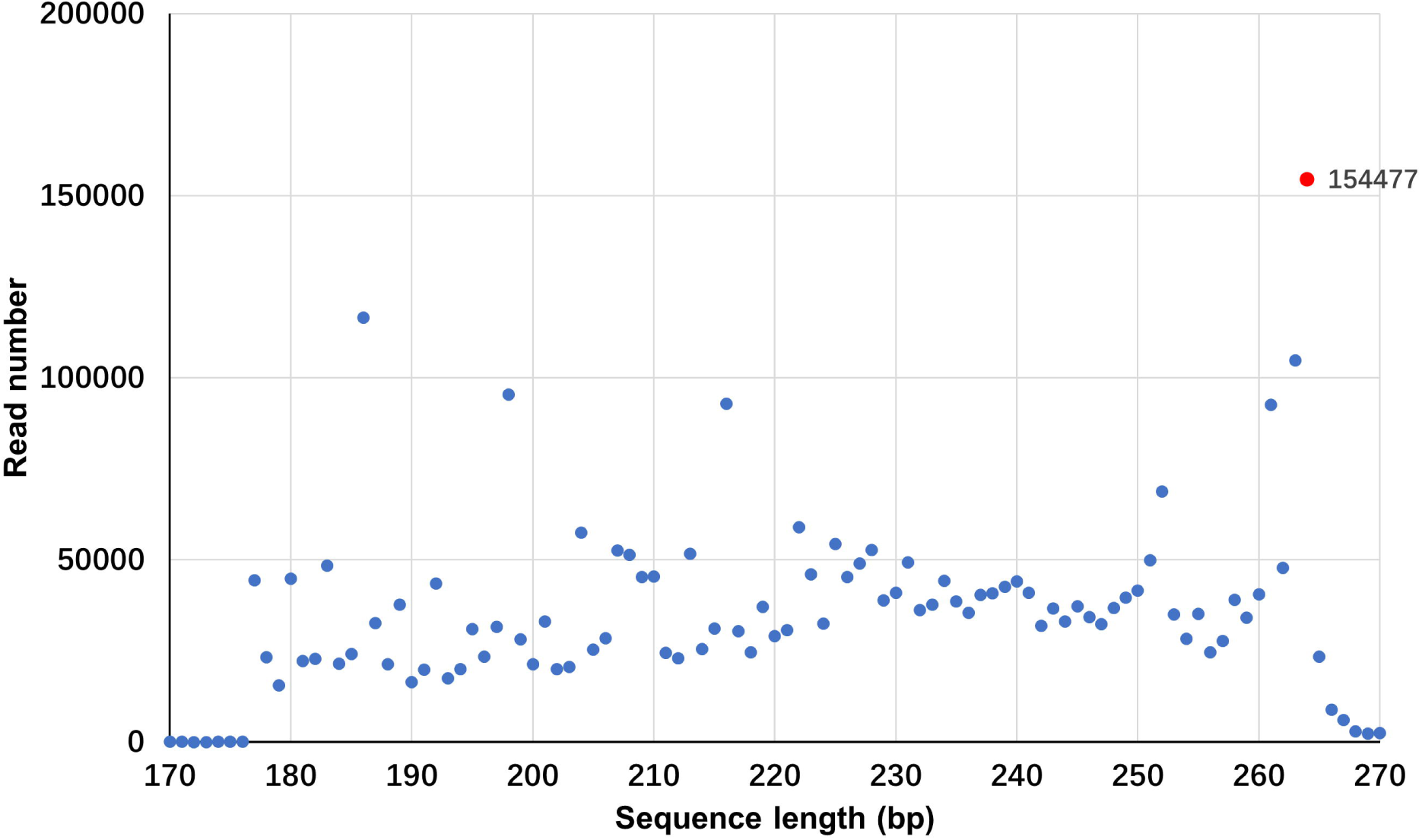
Read length distribution of HTS reads. Up- and down-stream ldsDNAs were cotransfected into HEK293T cells to form nanobody generating AND-gate genetic circuits. After RNA extraction, cDNA synthesis, PCR was employed to amplify nanobody CDR1-3 sequences. PCR products are subjected to pair-end HTS. Read length of merged clean reads containing both up- and down-stream ldsDNA sequence is summarized. The most abundant read length (264-bp) is marked in read.

### Abundance distribution of full nanobody ldsDNA sequences

In different display platforms, the capacity of the nanobody library usually serves as a determining factor of a successful experiment. The diversity of our nanobody library comes from 2 levels: the diversity of the input ldsDNAs and the combining diversity in the process of upstream- and downstream-ldsDNA linkage. In 154,477 reads of 264-bp full nanobody subset from all 3 biological repeats, we got 22,172 unique sequences. The two most abundant sequences have more than one thousand reads (1,117 and 1,044 reads respectively). There are 298 sequences containing 101 to 1,000 reads. And 19,737 sequences have only one read (Figure 3). Meanwhile, we investigated the redundancy among different biological repeats. As showed by the Venn diagram, only one sequence was present in all three independent nanobody libraries (Figure 4). Thus, this demonstrates that there is no obvious library redundancy when scaling up the experiments.

**Figure 3.**
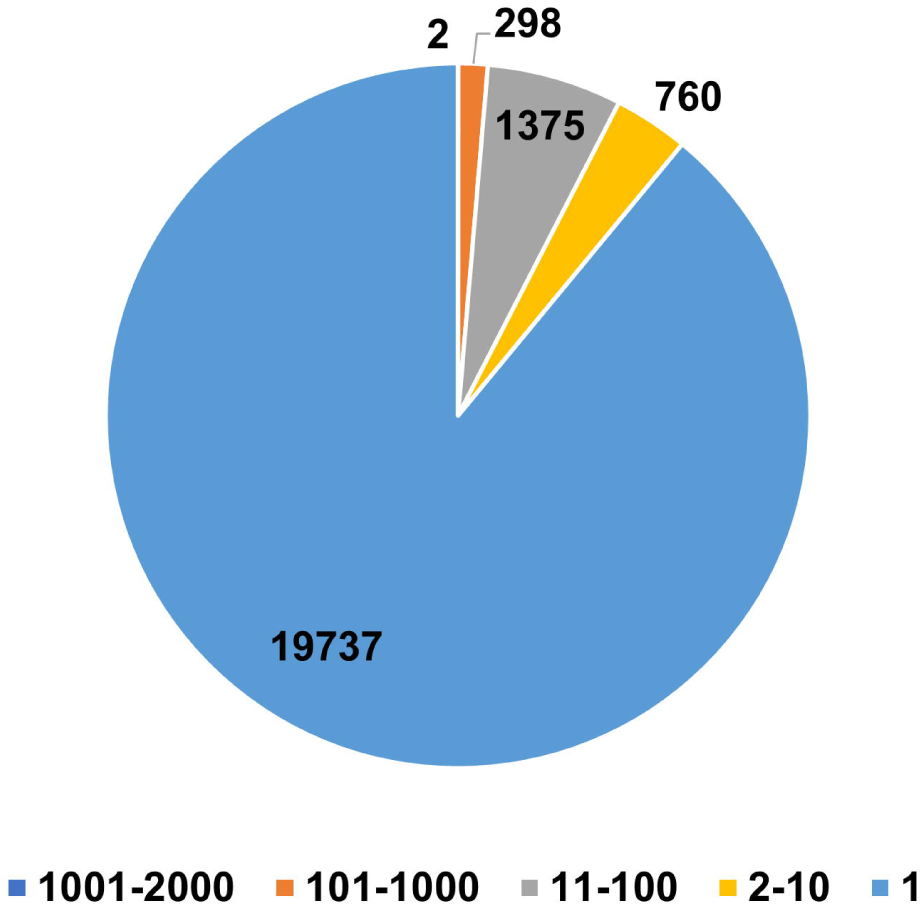
Abundance distribution of full nanobody ldsDNA sequences (264-bp). Sequences containing up- and down-stream ldsDNA sequences are included in analysis. Sequence abundance was divided into five groups: 1001-2000, 101-1000, 11-100, 2-10 and 1.

**Figure 4.**
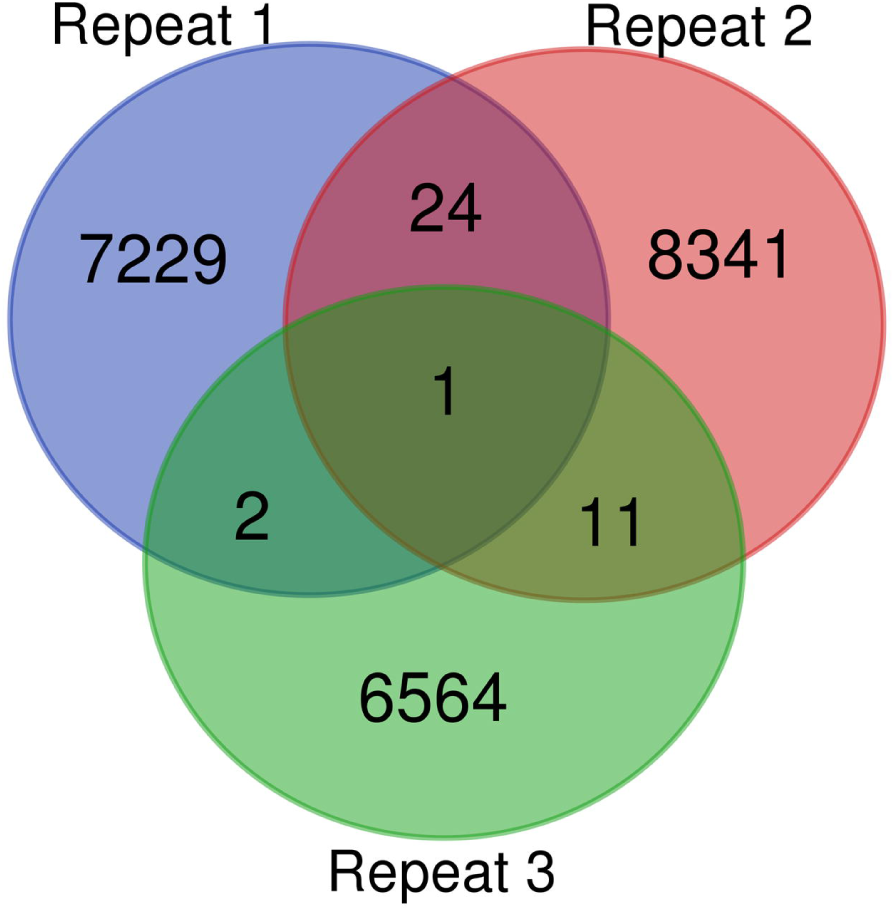
Venn diagram of unique sequence from three biological repeats. Full nanobody ldsDNA unique sequences (264-bp) are included in analysis.

### Nucleotide and amino acid distribution of nanobody library CDRs

We next analyzed the nucleotide distributions of CDRs in 22,172 unique 264-bp sequences. As shown in Figure 5A-C, nucleotide frequencies in each codon of the three CDRs are all in accordance with the NNK pattern. We also analyzed the composition of amino acid encoded by nanobody library CDR (Figure 5D). The STOP codon accounts for 2.64% in all CDR codons, which is lower than the theoretical frequency of the NNK degenerate codon (3.125%). The chance to get a pre-STOP codon increases with the elongation of the NNK peptide chain. When there are 24 NNK codons (8 in CDR1, 9 in CDR2, 7 in CDR3), the theoretical ratio of translation pre-termination is 46.7%. In our nanobody library, pre-STOP codon presents in 9,577 of the total 22,172 unique sequences (43.2%).

**Figure 5.**
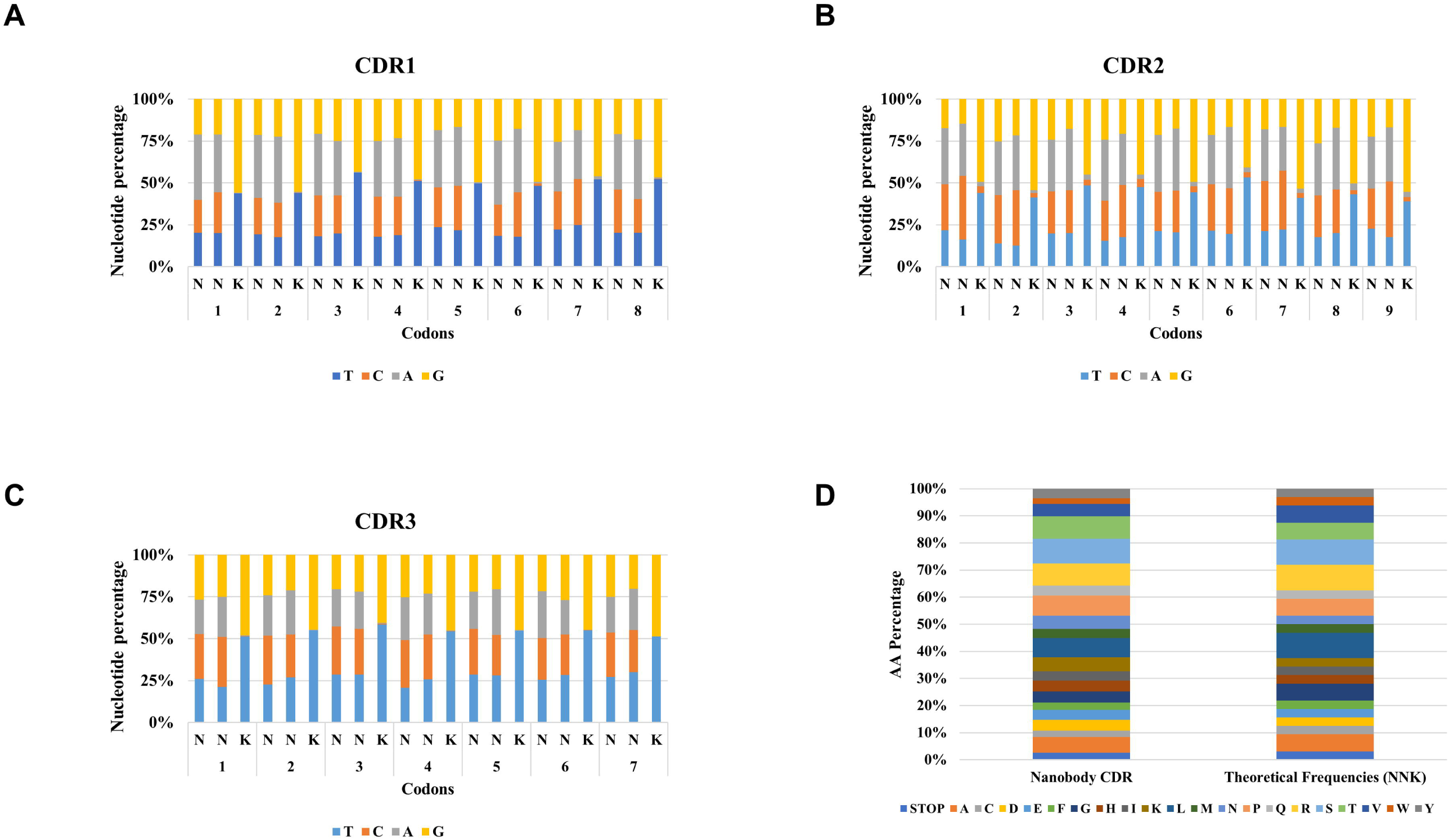
Nucleotide and amino acid distribution in nanobody library CDR codons. (A-C) Nucleotide frequencies in each codon of three CDRs are presented. The 22,172 unique full nanobody ldsDNA sequences (264-bp) were subjected to Perl script processing to extract oligonucleotide sequences corresponding to CDRs. The nucleotide composition of each codon was summarized and presented. (D) Observed and theoretical amino acid frequencies of nanobody CDRs. The CDR oligonucleotide sequences in (A-C) were translated into protein sequence. The amino acid composition of nanobody library was summarized and presented.

## Discussion

Here using **<u>l</u>**dsDNA-**<u>b</u>**ased **<u>A</u>**ND-**<u>g</u>**ate (LBAG) genetic circuit, we developed a novel method to construct nanobody library in mammalian cells. This protocol is easy to conduct, containing only two steps: PCR amplification and transfection of mammalian cell lines. Via PCR amplification, we introduce the CDRs into the upstream- and downstream-ldsDNAs respectively, which are then co-transfected into mammalian cell line to form AND-gate genetic circuits and launch the expression of full-length nanobodies. Through the linkage process, the diversity of the nanobody library is further increased. Through high throughput sequencing, we identified a total of 22,172 unique full-length sequences. And we further compared the library content among three independent biological repeats and found the non-redundant property of the nanobody library. These results demonstrate the feasibility of our strategy.

In this study, we employ GFP-nanobody as the framework of our library. Although GFP-nanobody has a high affinity for GFP (Kd = 0.32 nM), the length of its CDR3 (7AA) is much shorter than that of typical nanobodies (16-18AA) (1,5,33). Longer CDR3 is believed to contribute more antigen-binding conformation, which, to some extent, makes up for the decrease of binding capacity caused by the lack of light chain in nanobody. In future application, we can obtain nanobody library with longer CDR3 via utilizing longer PCR primers for downstream ldsDNAs. Another hallmark of nanobody is the interloop disulfide bond formed between CDR1 and CDR3. The disulfide bond stabilizes the domain and rigidifies the long CDR3 loop leading to stronger antigen interaction (38). By introducing Cysteine codons into both upstream- and downstream-ldsDNAs via primers, we could also establish interloop disulfide bonds in the nanobody library.

Here, our strategy is demonstrated to be feasible. However, there is still much room to improve on the efficiency of nanobody library construction. First, input ldsDNAs are frequently subjected to terminal nucleotide(s) deletion before their linkage. In our nanobody library, only 4.2% of the merged clean reads (154,477 / 3,679,989) contain the intact GFP-nanobody framework, although it is still the most abundant read length. Most of the other top read length are shorter than the intact framework, which is supposed to be the outcome of terminal nucleotide deletion (Figure 2). Terminal nucleotide(s) deletion of input ldsDNAs causes the loss of coding fragments and the destruction of nanobody framework, which leads to the shrinkage of the library size. One potential solution is to generate nuclease resistant ldsDNAs by introducing chemical modification into PCR products. We designed phosphorothioate-modified primers and introduced this modification into ldsDNAs via PCR amplification. However, we found that the presence of phosphorothioate bonds could block the formation of AND-gate genetic circuit (data not shown). Second, NNK degenerate codon brings biased amino acid composition and unwanted pre-stop and Cysteine codons. There is redundancy in both natural and NNK degenerate codon. For example, there are six and three Leucine coding codons in natural and NNK codes, respectively. For Phenylalanine, only one codon exists in both categories. This difference could lead to biased amino acid distribution (Figure 5D). The presents of pre-stop codon could also cut down the library size. In our nanobody library, 43.2% of 22,172 unique sequences contain pre-stop codon. And Cysteine could form disulfide bond with another Cysteine, which may interfere the tertiary structure of nanobody. We can synthesize primers by using trimer phosphoroamidites to get more balanced random CDRs meanwhile avoiding pre-stop and Cysteine codons (39,40).

In the present study, we only identified 22,172 unique nanobody coding sequences. This is not the exact library size generated by our strategy. First, only a small portion of extracted RNAs are reverse transcripted into cDNA. Second, not all cDNA is PCR amplified. Third, high throughput sequencing techniques could not sequence all PCR amplicon. So the library size we generated here should be larger than 22,172. However, we cannot give an estimate of the library size. Determining the nanobody sequence diversity in each cell by single-cell-sequencing will provide crucial information for estimating the whole library size.

In summary, we applied **<u>l</u>**dsDNA-**<u>b</u>**ased **<u>A</u>**ND-**<u>g</u>**ate (LBAG) genetic circuit to generate nanobody library, which may serve as a start point for further efforts to build high content nanobody library in mammalian cells.

## Acknowledgements

This work is supported in part by National Natural Science Foundation of China 31870860 (to S.L.), by grant “113” Plan for Talents of Innovation and Entrepreneurship in Xiqing District (to Tianjin Diagentech Co., Ltd), Tianjin. C.Z. is supported by Tianjin Science and Technology Support Key R & D Plan Project [19YFZCSY00420].

## Author Contributions

S.L. conceived and supervised the study. S.L. and C.Z. designed experiments. Y.Z. and C.Z. performed experiments. S.L., C.Z. and Y.Z. analyzed data, prepared figures, and wrote the paper. All authors discussed the results and reviewed the manuscript.

---

**Supplementary Table 1.**
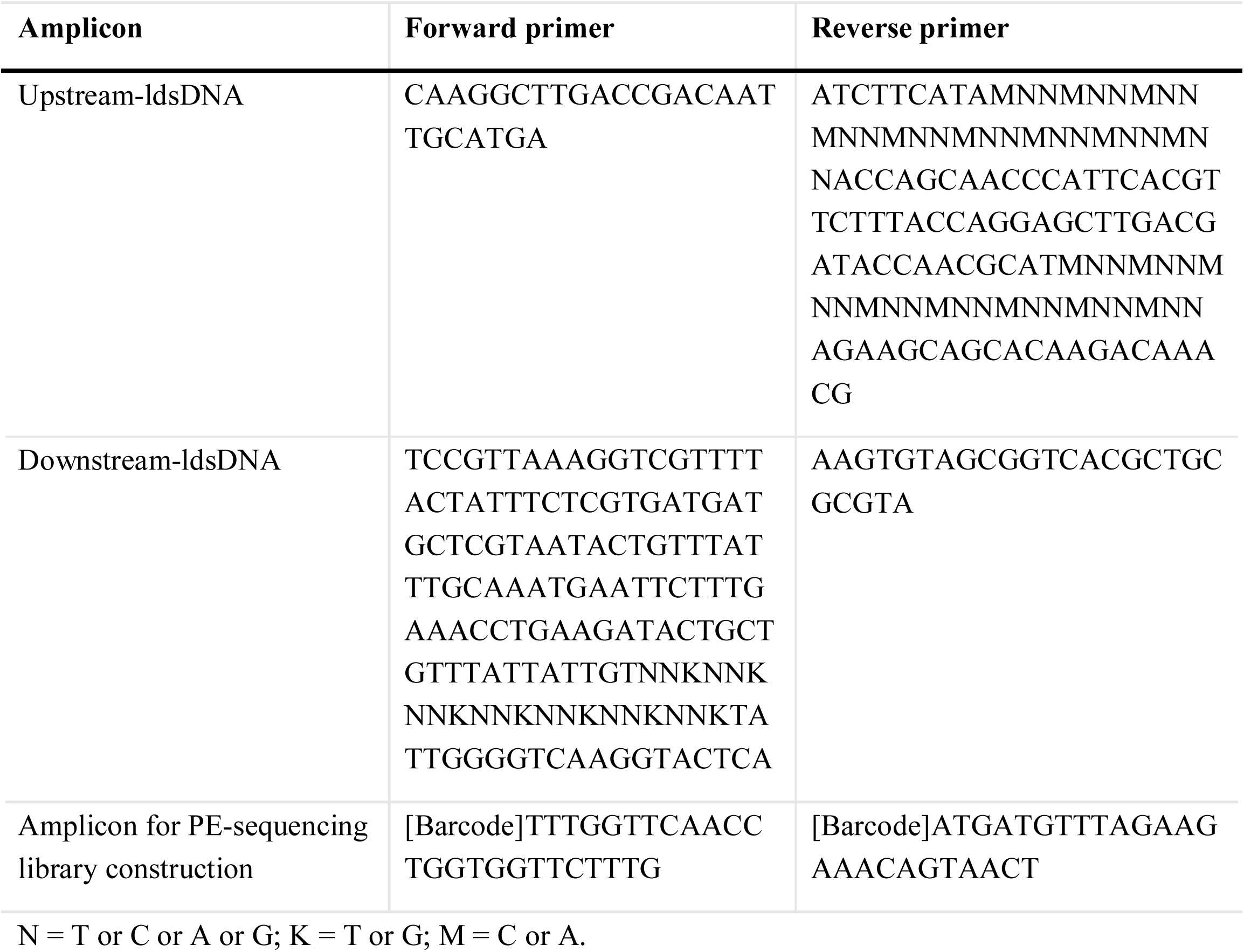
List of PCR primers used in study.

## Supplementary materials

Template ldsDNA sequences in study:

>Up-stream template ldsDNA

CAAGGCTTGACCGACAATTGCATGAAGAATCTGCTTAGGGTTAGGCGTTTTGCGCTGCTTCGCGATGTACGGGCCAGATATACGCGTTGACATTGATTATTGACTAGTTATTAATAGTAATCAATTACGGGGTCATTAGTTCATAGCCCATATATGGAGTTCCGCGTTACATAACTTACGGTAAATGGCCCGCCTGGCTGACCGCCCAACGACCCCCGCCCATTGACGTCAATAATGACGTATGTTCCCATAGTAACGCCAATAGGGACTTTCCATTGACGTCAATGGGTGGACTATTTACGGTAAACTGCCCACTTGGCAGTACATCAAGTGTATCATATGCCAAGTACGCCCCCTATTGACGTCAATGACGGTAAATGGCCCGCCTGGCATTATGCCCAGTACATGACCTTATGGGACTTTCCTACTTGGCAGTACATCTACGTATTAGTCATCGCTATTACCATGGTGATGCGGTTTTGGCAGTACATCAATGGGCGTGGATAGCGGTTTGACTCACGGGGATTTCCAAGTCTCCACCCCATTGACGTCAATGGGAGTTTGTTTTGGCACCAAAATCAACGGGACTTTCCAAAATGTCGTAACAACTCCGCCCCATTGACGCAAATGGGCGGTAGGCGTGTACGGTGGGAGGTCTATATAAGCAGAGCTCTCTGGCTAACTAGAGAACCCACTGCTTACTGGCTTATCGAAATTAATACGACTCACTATAGGGAGACCCAAGCTTGGTACCGCCACCATGGAGACAGACACACTCCTGCTATGGGTACTGCTGCTCTGGGTTCCAGGTTCCACTGGTGACTATCCATATGATGTTCCAGATTATGCTATGCAAGTTCAATTGGTTGAATCTGGTGGTGCTTTGGTTCAACCTGGTGGTTCTTTGCGTTTGTCTTGTGCTGCTTCT

> Down-stream template ldsDNA

TATTGGGGTCAAGGTACTCAAGTTACTGTTTCTTCTAAACATCATCATCATCATCATGAACAAAAACTCATCTCAGAAGAGGATCTGAATGCTGTGGGCCAGGACACGCAGGAGGTCATCGTGGTGCCACACTCCTTGCCCTTTAAGGTGGTGGTGATCTCAGCCATCCTGGCCCTGGTGGTGCTCACCATCATCTCCCTTATCATCCTCATCATGCTTTGGCAGAAGAAGCCACGTTAGTCTAGAGGGCCCTATTCTATAGTGTCACCTAAATGCTAGAGCTCGCTGATCAGCCTCGACTGTGCCTTCTAGTTGCCAGCCATCTGTTGTTTGCCCCTCCCCCGTGCCTTCCTTGACCCTGGAAGGTGCCACTCCCACTGTCCTTTCCTAATAAAATGAGGAAATTGCATCGCATTGTCTGAGTAGGTGTCATTCTATTCTGGGGGGTGGGGTGGGGCAGGACAGCAAGGGGGAGGATTGGGAAGACAATAGCAGGCATGCTGGGGATGCGGTGGGCTCTATGGCTTCTGAGGCGGAAAGAACCAGCTGGGGCTCTAGGGGGTATCCCCACGCGCCCTGTAGCGGCGCATTAAGCGCGGCGGGTGTGGTGGTTACGCGCAGCGTGACCGCTACACTT

>GFP-nanobody AA sequence (CDR1-CDR2-CDR3)

QVQLVESGGALVQPGGSLRLSCAASGFPVNRYSMRWYRQAPGKEREWVAGMSSAGDRSSYEDSVKGRFTISRDDARNTVYLQMNSLKPEDTAVYYCNVNVGFEYWGQGTQVTVSS

>264-bp sequence (CDR1-CDR2-CDR3)

TCTTGTGCTGCTTCTNNKNNKNNKNNKNNKNNKNNKNNKATGCGTTGGTATCGTCAAGCTCCTGGTAAAGAACGTGAATGGGTTGCTGGTNNKNNKNNKNNKNNKNNKNNKNNKNNKT ATGAAGATTCCGTTAAAGGTCGTTTTACTATTTCTCGTGATGATGCTCGTAATACTGTTTA TTTGCAAATGAATTCTTTGAAACCTGAAGATACTGCTGTTTATTATTGTNNKNNKNNKNN KNNKNNKNNKTATTGGGGTCAAGGT

